# Dental restoration longevity among geriatric and adult special needs patients

**DOI:** 10.1101/202069

**Authors:** D. J. Caplan, Y. Li, W. Wang, S. Kang, L. Marchini, H. J. Cowen, J. Yan

## Abstract

This study aimed to describe the survival trajectory of dental restorations placed in an outpatient population of geriatric and adult special needs patients over a 15-year span, with particular interest in longevity of subsequent restorations in teeth that received multiple restorations over time. Dental restorations of different types and sizes in patients age ≥65 years treated between 2000-14 at the University of Iowa, College of Dentistry were followed until they incurred an event (i.e., restoration replacement, extraction of the tooth, or endodontic treatment of the tooth). Survival analysis and extended Cox regression models were used to generate hazards ratios for selected predictor variables. A total of 9184 restorations were followed in 1551 unique patients. During the follow-up period, 28.7% of these restorations incurred an event; and overall the restorations had a median lifespan of 6.25 years. In multivariable regression models, after controlling for gender and age, composite restorations and greater number of restoration surfaces were associated with higher risks of failure; and the initial restoration recorded in the database for each subject tended to have lower risk of failure than restorations placed later that included any of those same surfaces. This information potentially could be helpful to elderly patients considering various restorative treatment options during the dental treatment planning and informed consent process.

## Background

The American elderly population is expected to grow from 43.1 million in 2012 to 83.7 million by the year 2050 – a 94.2% increase (Ortman et al. 2014). The growing population of elderly is having a pronounced impact on society and an even more dramatic influence on the U.S. health care system (WHO 2015). Elderly people have more chronic diseases and consequently bear a disproportionate share of the global burden of disease (Prince et al. 2015).

Disease and disability that may come with age have been linked to poor oral health (Zhang et al. 2017), which in turn can play a role in life-threatening systemic health complications (van der Maarel-Wierink et al. 2013; Beikler and Flemmig 2011). Caries in permanent teeth was the most prevalent health condition worldwide in 2010 (Kassebaum et al. 2015) and is one of the most prevalent conditions among the elderly population as well, mainly among those who are frail (Thomson 2014).

Cavitated dental lesions are commonly treated with different materials and restoration types, and the longevity of these restorations has been the focus of much research and debate (Astvaldsdottir et al. 2015; Burke and Lucarotti 2009; Demarco et al. 2015; Demarco et al. 2012; Dobloug and Grytten 2015; Forss and Widstrom 2004; Kolker et al. 2005; Laske et al. 2016; Moraschini et al. 2015; Opdam et al. 2011; Schwendicke et al. 2016), yet the majority of these studies have evaluated restorations placed in the general population, so most published studies do not account for the challenges faced by the frail elderly. These challenges are numerous and include systemic health, oral health, and social factors. Systemic health factors commonly include dementia (Brennan and Strauss 2014), depression (Hybels et al. 2016), diabetes (de Deco et al. 2007), stroke (Lam et al. 2013), arthritis (Tavares et al. 2014), and polypharmacy (Singh and Papas 2014), among others. Oral health factors include wearing removable dentures, presenting with a heavily restored dentition, or having poor oral hygiene, gingival recession with root exposure, or xerostomia (Jablonski and Barber 2015; Singh and Papas 2014). Social factors include financial limitation, dependence on caregivers, institutionalization, ageism, and access to appropriate care (Friedman et al. 2014).

Considering the lack of studies that address the longevity of dental restorations placed among the elderly population, the aims of this paper were to 1) describe the survival trajectory of restorations placed in an academic geriatric and special needs outpatient clinic over a period of 15 years, and 2) evaluate factors that potentially influence restoration longevity in this traditionally underserved population. Of particular interest was to quantify the degree to which subsequent, additional restorative therapy on a given tooth was related to restoration longevity. Our hypothesis was that controlling for all available patient-and restoration-level variables, the initial restoration recorded in the database would experience greater longevity than would subsequent restorations.

## Methods

Since the mid-1980’s the Geriatric and Special Needs (GSN) Dentistry Clinic at the University of Iowa, College of Dentistry (COD) has offered comprehensive dental care to geriatric patients, those on numerous medications, and adults with behavioral, psychological, or other health conditions. Care is provided by faculty-supervised senior dental students and graduate students. Consent prior to dental treatment is obtained from the patients or their health care powers-of-attorney.

Prior to conducting this study, Institutional Review Boards at both the University of Iowa and the University of Connecticut independently declared the project not to require their regulatory oversight because no personal health information was included in the working dataset.

For this analysis, electronic data were obtained for all dental procedures delivered during the 15-year period from 1/1/00 - 12/31/14. Initially we identified all patients of at least 65 years of age who had at least one American Dental Association (ADA) procedure code representing an intracoronal or extracoronal restoration placed in the GSN clinic during the 15-year period. The specific ADA restorative codes were D2140-D2161 for amalgam restorations; D2330-D2335 and D2385-D2394 for composite restorations; D2330.1-D2335.1 and D2385.1-D2394.1 for glass ionomer cement (GIC) restorations (the “.1” is a COD designation to indicate GIC as opposed to composite); D2740-D2752 and D2790-D2792 for crowns; and D6750-D6792 for bridges. Associated with each restorative procedure was the following information: Tooth number, surfaces (mesial, distal, lingual, facial / buccal, incisal / occlusal); number of surfaces restored (1,2,3+); restoration type (amalgam, composite, GIC, crown, bridge retainer); cohort (2000-04, 2005-09, 2010-14); provider type (pre-doctoral student versus graduate student / faculty); payment method (Medicaid, self-pay, private insurance); and restoration sequence (first restoration captured in the database, or “index restoration”, versus subsequent restoration). The patient’s age at the time of each procedure, as well as sex, also were available electronically.

We then identified all procedures of any type received by these patients at any COD clinic after the index restoration placement date. Restorations placed on the patient’s last visit to the COD were not eligible for follow-up. Restorations that were followed included amalgam, composite, and GIC restorations (placed intracoronally, in specific surfaces within a tooth), plus crowns and bridge retainers (placed extracoronally, covering all coronal surfaces of a tooth). For intracoronal restorations that were followed, any subsequent intracoronal restoration of any of the same surfaces indicated an event, as did subsequent placement of a crown or bridge retainer on that tooth. For extracoronal restorations that were followed, any subsequent restoration placed on any surface indicated an event. For all restorations followed, extraction or endodontic therapy of the tooth also indicated an event for the restoration.

For all restorations follow-up began on the date of restoration placement. For restorations that incurred an event, the event date marked the end of follow-up. Restorations that did not incur an event throughout the entire follow-up period were censored on the date of the patient’s last visit to any COD clinic for any reason, thus each restoration that was followed either incurred an event or was censored.

Because each tooth could have had multiple restorations eligible for follow-up, and because each subsequent restoration could be considered as both a failure for the preceding restoration and a new restoration that could be followed in its own right, statistical analyses reflected the correlated nature of the observations. All data analyses were conducted using R (R_Core_Team 2017). Univariate analyses were conducted to assess distributions and to perform range checks of all the variables. Frequency tables of single variables and contingency tables of combinations of variables also were generated to assess patterns and dependence structures, and survival curves were estimated by the Kaplan-Meier estimator (Kaplan and Meier 1958). Kaplan-Meier survival curve estimates were obtained from R package “prodlim” (Gerds 2017), which accounts for within-patient dependence (Ying and Wei 1994). Based on those procedures we defined and selected the subset of variables to be employed in regression models.

To account for within-patient dependence, a Cox proportional hazards model with a patient level random effect was fitted (Therneau 2015; Therneau and Grambsch 2000). This model is an extended Cox model (Cox 1972) where the hazard of an individual restoration depends not only on the predictor variables and a baseline hazard but also on an unobserved patient-level random effect (also known as frailty). This approach has been used previously in dental research in modeling clustered event times to account for within-patient dependence (Kopperud et al. 2012; Opdam et al. 2014). The model is specified by

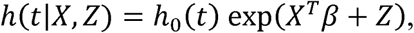

where *h*(*t*|*X, Z*) is the hazard function of a restoration in a patient with predictor vector X and unobserved frailty Z, and *h*_0_(*t*) is an unspecified baseline hazard function. The baseline hazard could differ for each combination of values for stratified variables.

We used the R package “survival” (Therneau 2015) to fit the frailty Cox model and verify the proportional hazards assumption (Grambsch and Therneau 1994). If the effect of a predictor did not satisfy the proportional hazards assumption we used two remedies. The first was to stratify on this variable (if it was categorical) so that each stratum had its own baseline hazard function. The second was to allow the effect of this predictor to change over time in a piecewise constant fashion.

## Results

A total of 9184 restorations were followed, including 4670 (50.8%) in female patients; and 1551 unique subjects, 835 (53.8%) of whom were female (data not shown). The median number of followed restorations per subject was four; and 1615 out of 6841 teeth (23.6%) had more than one restoration (data not shown).

Table 1 describes the numbers and types of restorations that were followed in this analysis. Amalgam, composite, and GIC each comprised about one-third of the restorations, with only a small percentage of crowns or bridge retainers placed. The overall censoring rate was 71.3%, with crowns and bridges having higher censoring rates (~84%) than any of the intracoronal restorations (~71%). In general, intracoronal restorations were most often replaced with the same type of restoration placed originally. Of the intracoronal restorations, amalgam and GIC restorations were more likely to fail due to tooth extraction than were composite restorations, and bridges were the most common type of restoration to incur an event due to endodontic therapy.

**Table 1:** Types of Restorations and Failures

| **Type of Failure** | **Type of Restoration (all followed restorations)** |  |  |  |  | **Total** |
| --- | --- | --- | --- | --- | --- | --- |
|  | **Amalgam** | **Composite** | **GIC** | **Crown** | **Bridge** |  |
| TOTAL (N) | 3011 | 2565 | 3295 | 236 | 77 | 9184 |
| Censored<br>(no failure) | 71.2% | 71.3% | 70.2% | 84.7% | 84.4% | 71.3% |
| Amalgam | 6.9% | 3.0% | 2.6% | 2.1% | 3.9% | 4.1% |
| Composite | 3.8% | 11.4% | 3.6% | 0.4% | 0.0% | 5.7% |
| Glass Ionomer | 3.9% | 5.2% | 10.1% | 2.5% | 1.3% | 6.4% |
| Crown | 1.6% | 0.8% | 0.3% | 1.7% | 0.0% | 0.9% |
| Bridge | 0.3% | 0.2% | 0.1% | 0.4% | 0.0% | 0.2% |
| Extraction | 11.6% | 7.2% | 12.6% | 7.6% | 5.2% | 10.6% |
| Endodontic Therapy | 0.7% | 0.9% | 0.6% | 0.4% | 5.2% | 0.7% |

| **Type of Failure** | **Type of Restoration (only failed restorations)** |  |  |  |  | **Total** |
| --- | --- | --- | --- | --- | --- | --- |
|  | **Amalgam** | **Composite** | **GIC** | **Crown** | **Bridge** |  |
| TOTAL (N) | 866 | 736 | 983 | 36 | 12 | 2633 |
| Amalgam | 24.0% | 10.6% | 8.9% | 13.9% | 25.0% | 14.5% |
| Composite | 13.3% | 39.8% | 11.9% | 2.8% | 0.0% | 20.0% |
| Glass Ionomer | 13.4% | 18.2% | 33.9% | 16.7% | 8.3% | 22.4% |
| Crown | 5.4% | 2.9% | 1.0% | 11.1% | 0.0% | 3.1% |
| Bridge | 1.0% | 0.5% | 0.2% | 2.8% | 0.0% | 0.6% |
| Extraction | 40.3% | 25.0% | 42.2% | 50.0% | 33.3% | 36.8% |
| Endodontic Therapy | 2.5% | 3.0% | 1.9% | 2.8% | 33.3% | 2.6% |

Figures 1A-G show Kaplan-Meier survival curves (with 95% confidence intervals) for the intracoronal restorations: overall (Figure 1A), by restorative material across tooth types (Figures 1B-D), and by restorative material across number of restored surfaces (Figures 1E-G). These curves illustrate the relative survival times not considering the effect of potential confounding variables. The median lifespan for all intracoronal restorations was 6.25 years (Figure 1A). Figures 1B-D show that the median lifespans for amalgam, composite and GIC restorations, respectively, in anterior teeth were 8.09, 5.08 and 4.28 years; in premolars they were 8.00, 6.39 and 4.88 years; and in molars they were 6.99, 6.38 and 4.66 years. For amalgam restorations, median survival times were similar for anterior teeth and premolars but worse for molars; for composite restorations, they were worse for anterior teeth than for premolars and molars; and for GIC restorations, they were similar across all tooth types. Regardless of restorative material, larger restorations tended to have shorter lifespans than smaller restorations (Figures 1E-G).

**Figure 1A-G:**
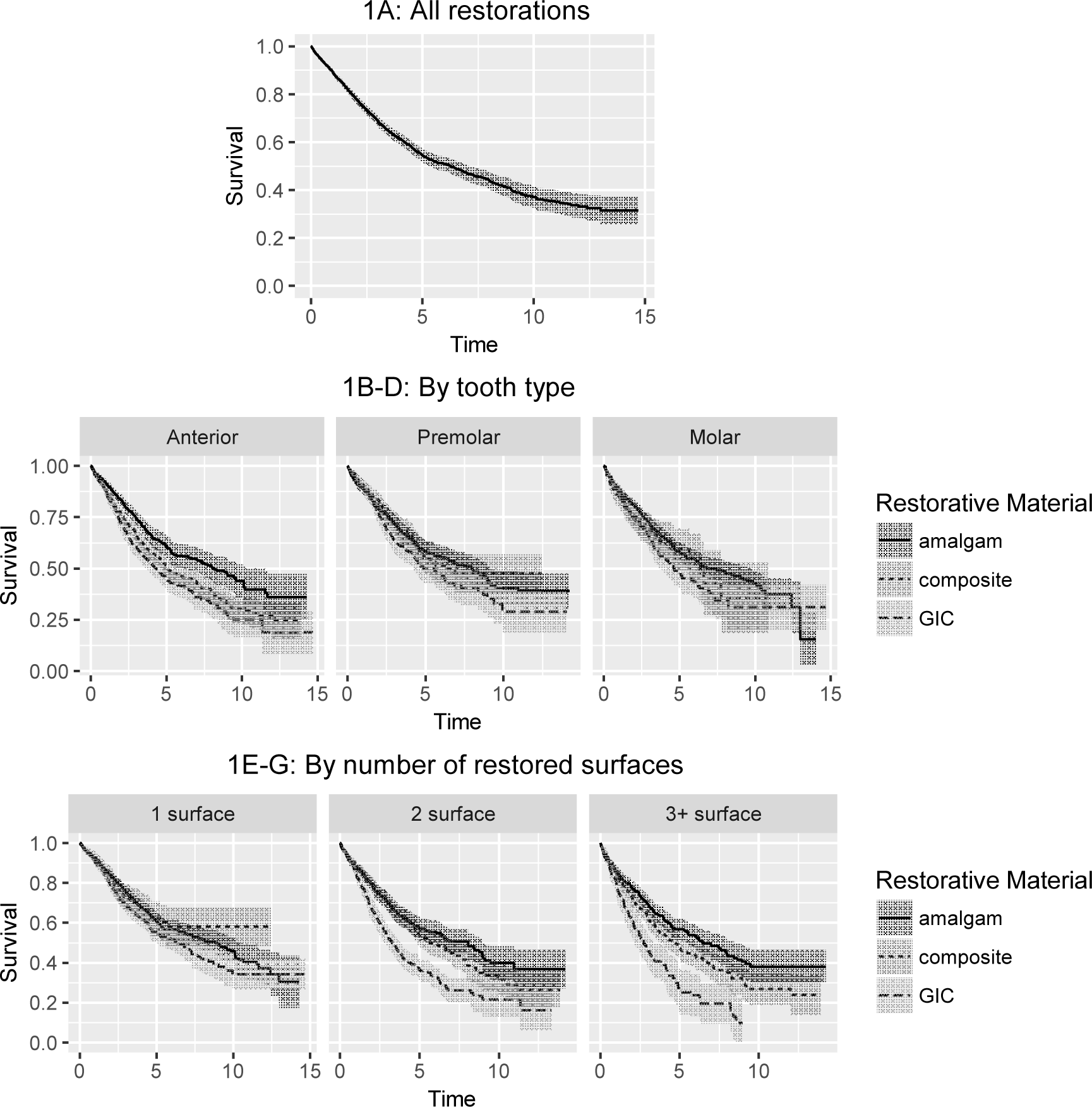
Kaplan-Meier survival curves for all (A), amalgam, composite, and glass ionomer restorations by tooth type (B-D) and by number of restored surface (E-G)

Table 2 presents median survival times from the Kaplan-Meier curves and hazard ratios (HRs) from the Cox models for all patients and different subgroups of patients. Subgroups represented strata based on tooth type and restorative material. For each subgroup, estimated HRs and their 95% confidence intervals for two variables are reported after controlling for sex, age, cohort, provider type and payment method: 1) number of restored surfaces; and 2) whether or not the restoration was the second or later restoration in the recorded data for a given tooth. The reported models all satisfied the proportional hazards assumption. Patterns repeated consistently across regression models were: a) restorations with more restored surfaces tended to fail sooner (e.g., compared to 1-surface restorations, statistically significant HRs of 1.39 for 2-surface restorations and 1.76 for larger restorations were observed); and b) restorations that were the second or later restoration in the recorded data for a given tooth tended to fail sooner (e.g., compared to index restorations, statistically significant HRs ranged from 1.20 – 1.89, depending on tooth type and restorative material).

**Table 2:** Median Survival Times and Hazard Ratios* for Different Tooth Types and Restorative Materials

| Among | Restorative Material | Median Survival Time (Years) | Number of Restored Surfaces^ | Second or Later Restoration |
| --- | --- | --- | --- | --- |
| **All** | **All** | 6.25 | M = 1.39 (1.25 - 1.54)<br>L = 1.76 (1.57 - 1.97) | 1.26 (1.10 - 1.45) |
|  | **Amalgam** | 7.95 | M = 1.31 (1.05 - 1.62)<br>L = 1.51 (1.21 - 1.89) | 1.46 (1.25 - 1.71) |
|  | **Composite** | 5.49 | M = 1.42 (1.01 - 2.00)<br>L = 1.87 (1.32 - 2.65) | 1.32 (1.11 - 1.57) |
|  | **Glass Ionomer** | 4.56 | M = 1.64 (1.40 - 1.93)<br>L = 2.24 (1.81 - 2.76) | 1.27 (1.09 - 1.48) |
| **Molars** | **All** | 6.23 | M = 1.42 (1.14 - 1.76)<br>L = 1.68 (1.31 - 2.15) | 1.44 (1.18 - 1.75) |
|  | **Amalgam** | 6.99 | M = not significant<br>L = 1.55 (1.16 - 2.09) | 1.32 (1.01 - 1.73) |
|  | **Composite** | 6.38 | M = not significant<br>L = not significant | 1.82 (1.04 - 3.20) |
|  | **Glass Ionomer** | 4.66 | M = 2.06 (1.42 - 2.99)<br>L = 2.87 (1.58 - 5.19) | 1.78 (1.24 - 2.55) |
| **Premolars** | **All** | 6.43 | M = 1.57 (1.26 - 1.97)<br>L = 1.53 (1.19 - 1.97) | 1.29 (1.06 - 1.57) |
|  | **Amalgam** | 8.00 | M = not significant<br>L = not significant | Not significant |
|  | **Composite** | 6.39 | M = 1.75 (1.11 - 2.76)<br>L = not significant | 1.52 (1.01 - 2.30) |
|  | **Glass Ionomer** | 4.88 | M = 2.01 (1.38 - 2.93)<br>L = 2.81 (1.77 - 4.47) | 1.51 (1.09 - 2.10) |
| **Anterior** | **All** | 5.19 | M = 1.36 (1.13 - 1.64)<br>L = 1.85 (1.51 - 2.26) | Not significant |
|  | **Amalgam** | 8.09 | Not included in model due to small sample size | 1.62 (1.21 - 2.17) |
|  | **Composite** | 5.08 | Not included in model due to stratification | 1.27 (1.04 - 1.55) |
|  | **Glass Ionomer** | 4.28 | M = 1.36 (1.09 - 1.70)<br>L = 1.79 (1.36 - 2.37) | not significant |

\*Controlling for sex, age, cohort, provider type and payment method
^M = “Medium” (restorations with 2 surfaces); L = “Large” (restorations with 3+ surfaces)

## Discussion

In the present sample of restorations placed in an outpatient geriatric and adult special needs population, two factors were consistently associated with shorter restoration longevity, controlling for all available variables: larger restorations and restorations placed later chronologically within a given tooth.

Larger restorations have been associated with shorter lifespan in previously published studies (Demarco et al. 2012; Laske et al. 2016). In one of these, multi-surface restorations failed more readily than single surface restorations, and every surface added to a restoration increased the risk of failure by 40% (Demarco et al. 2012). Among geriatric and adult special needs populations this finding is particularly important, as there is a tendency for these patients to present with a more heavily restored dentition and generally larger restorations (Jablonski and Barber 2015).

Our finding that chronologically earlier restorations within teeth had longer survival than subsequent ones is a novel finding, but is intuitive in some regards. All other things being equal, subsequent intracoronal restorations (even if identical surfaces are restored) likely remove more tooth structure than did the initial restoration, which potentially could lead to a greater rate of failure due to fracture -- though most often, authors that analyze “restoration size” typically are referring to the number of surfaces restored (Burke and Lucarotti 2009; Demarco et al. 2012; Laske et al. 2016) rather than restoration volume. As reported previously, restorations with three or more surfaces present with a relative risk of failure of 3.3 when compared to single-surface restorations (Demarco et al. 2012).

Some factors previously reported as associated with restoration longevity in previous studies, such as restorative material (Burke and Lucarotti 2009; Moraschini et al. 2015; Schwendicke et al. 2016) and tooth type (Demarco et al. 2012; Laske et al. 2016) were not consistently associated with longevity in the present study. A general population likely would have more opposing teeth than the present sample of elderly patients (though opposing dentition could not be assessed using the present study design), and opposing teeth would be expected to confer a greater opportunity for failure due to fracture of teeth or restorations. Occlusal restorations in teeth of general or younger populations likely would experience higher masticatory forces as well, and if so, those teeth and restorations likely would be more susceptible to failure. In this geriatric and special needs population, impaired masticatory forces associated with systemic (Tavares et al. 2014) or oral health factors (Thomson 2014) might reduce the differences in failure across tooth types.

Like other retrospective analyses using existing electronic databases, this study was limited to available, consistently-recorded variables, any of which could contain errors. Another limitation of this study is that what we called the “index restoration” was not necessarily the first restoration in a tooth, but rather the first that appeared chronologically in the database. Nevertheless, to our knowledge this study is the first to assess restoration longevity in a geriatric and adult special needs population for as long as 15 years, including data from multiple restorations within teeth and multiple events over time. Future prospective studies among this population are warranted so that elderly patients can be better informed about their restorative treatment options.

## Conclusion

Restorations placed in a geriatric and adult special needs outpatient dental clinic had a median lifespan of 6.25 years. Multivariable regression models showed that the greater the number of restoration surfaces, the shorter the restoration longevity; and that restorations appearing chronologically as first in the database lasted longer compared to subsequent restorations within a given tooth. To our knowledge, this information is the first of its kind about restoration longevity in elderly adult populations, and should be useful to both patients and restorative dentists during the dental treatment planning and informed consent process.

## Author Contributions

This project was directed by DJC and primarily conceived by DJC, JY, and SK. DJC was responsible for data acquisition, JY and SK contributed to the study design, and JY directed the statistical analysis. HJC supervised restoration placement and provided additional clinical oversight for many of the restorations analyzed. YL and WW developed R scripts for data preparation, data analysis, and statistical report. LM contributed to result interpretation and conceptual aspects of the work. All authors critically revised numerous drafts of the manuscript and gave final approval for its submission.

